## Supplementary Materials for "Human brain state dynamics reflect individual neuro-phenotypes"

### Supplementary Figures

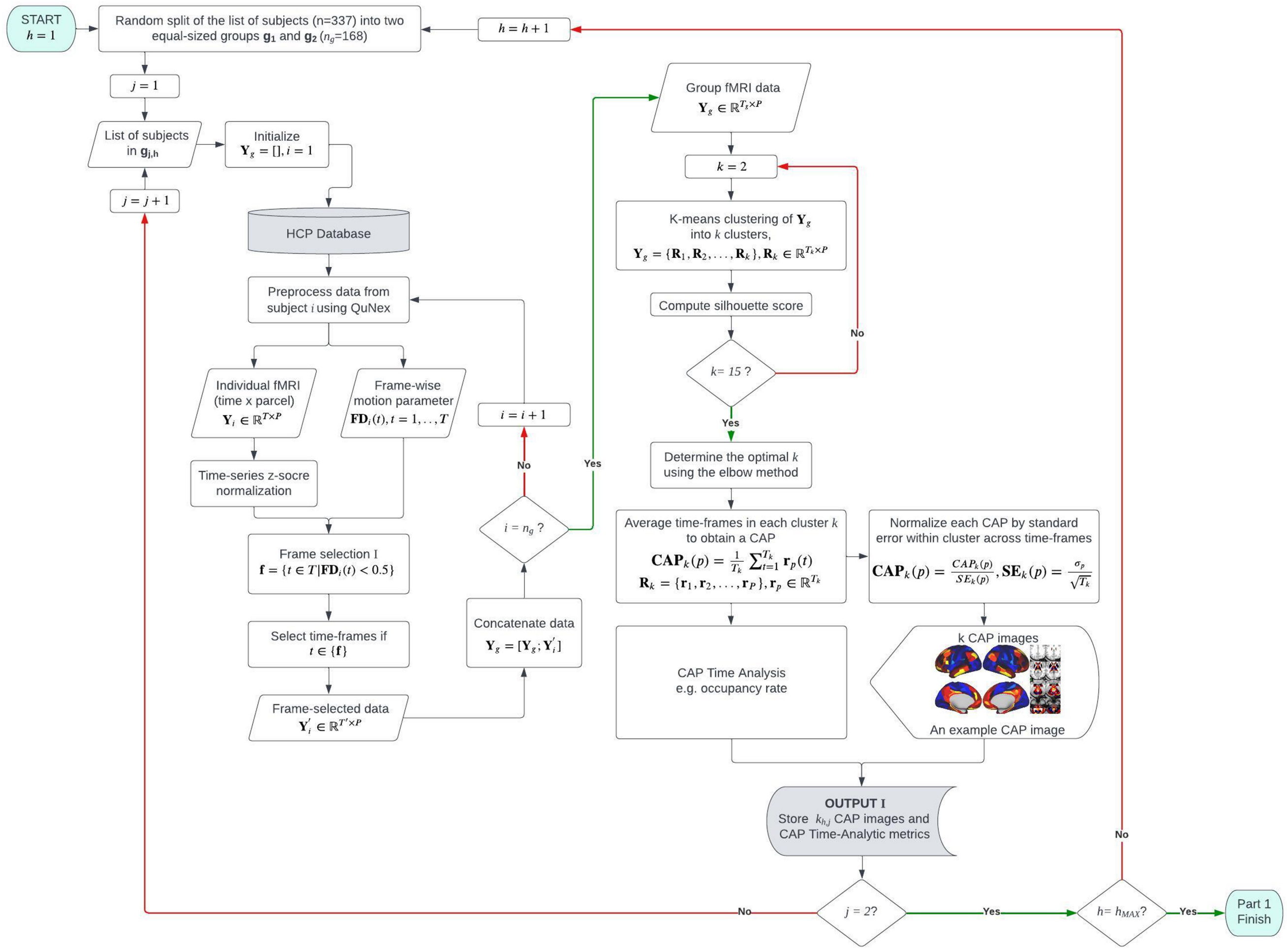


**Fig. S1. Workflow of CAP analysis.**

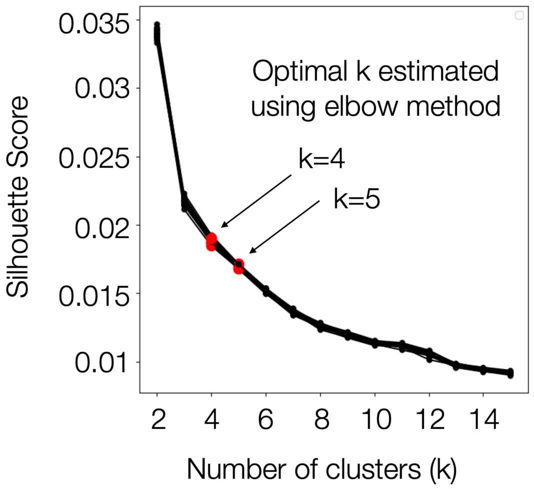


**Fig. S2. Quality of the K means clustering solution in CAP analysis.** Silhouette scores were estimated across different numbers of clusters ($k$) from the K-means clustering solution from a split data. Results from 10 permutations (two split-halves in each permutation) are shown. Optimal $k$ values were estimated using the elbow method for the Silhouette scores and are highlighted in red.

DRAFT

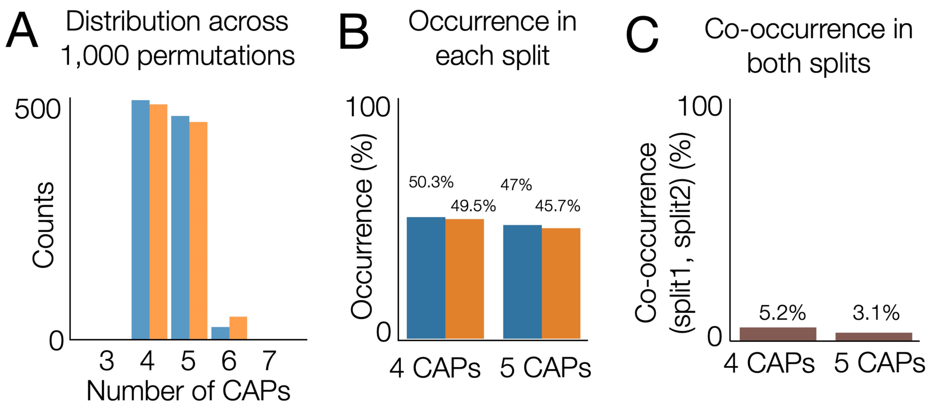


**Fig. S3. Occurrence of CAPs across permutations.** (A) The estimated number of CAPs ($k$) in each split across 1,000 permutations. (B) Occurrence rate ($\%$) of k=4 or k=5 solutions in each split. (C) Co-occurrence rate ($\%$) of $k = 4$ or $k = 5$ solutions in both splits.

DRAFT

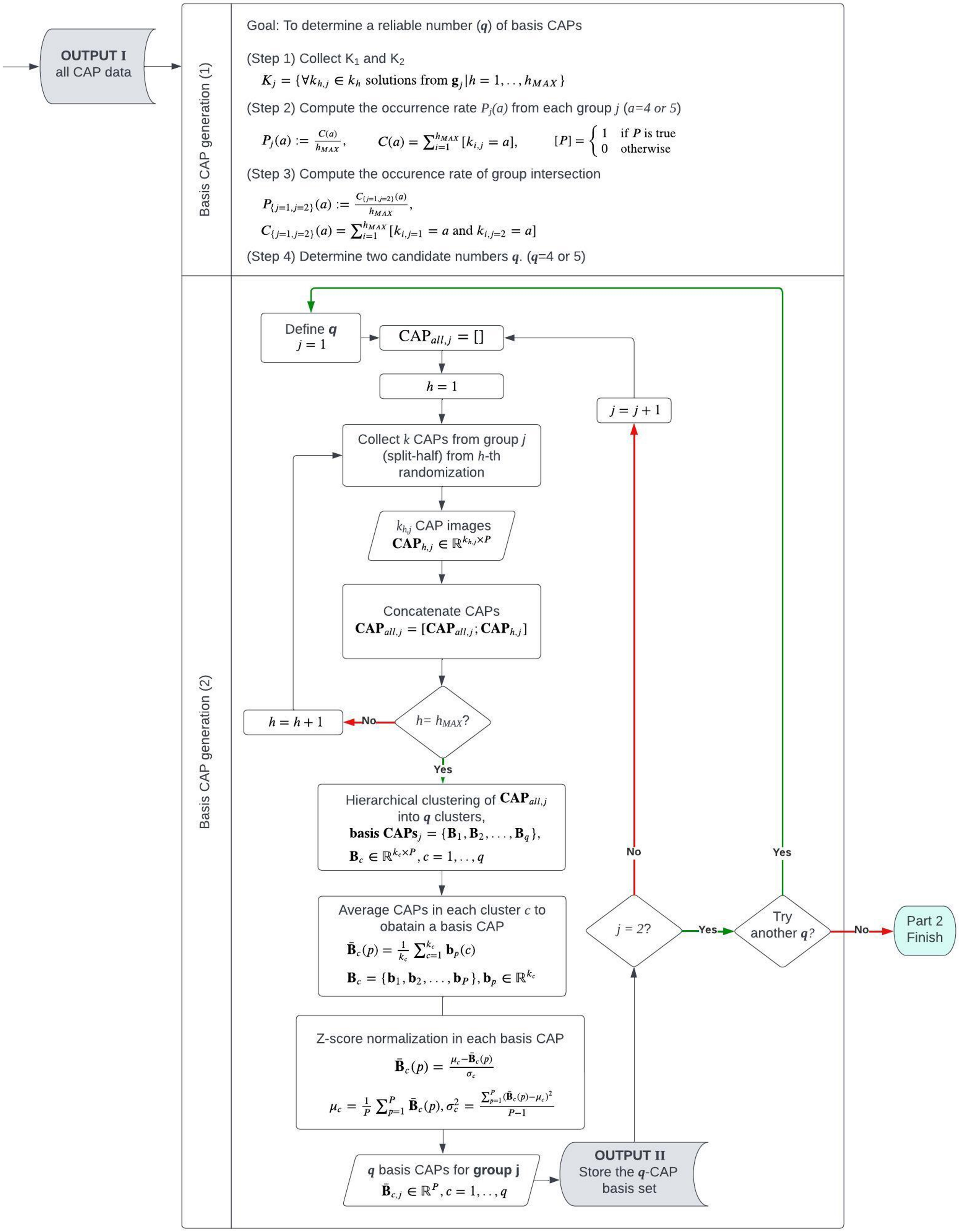


**Fig. S4. Generation of basis CAP sets.**

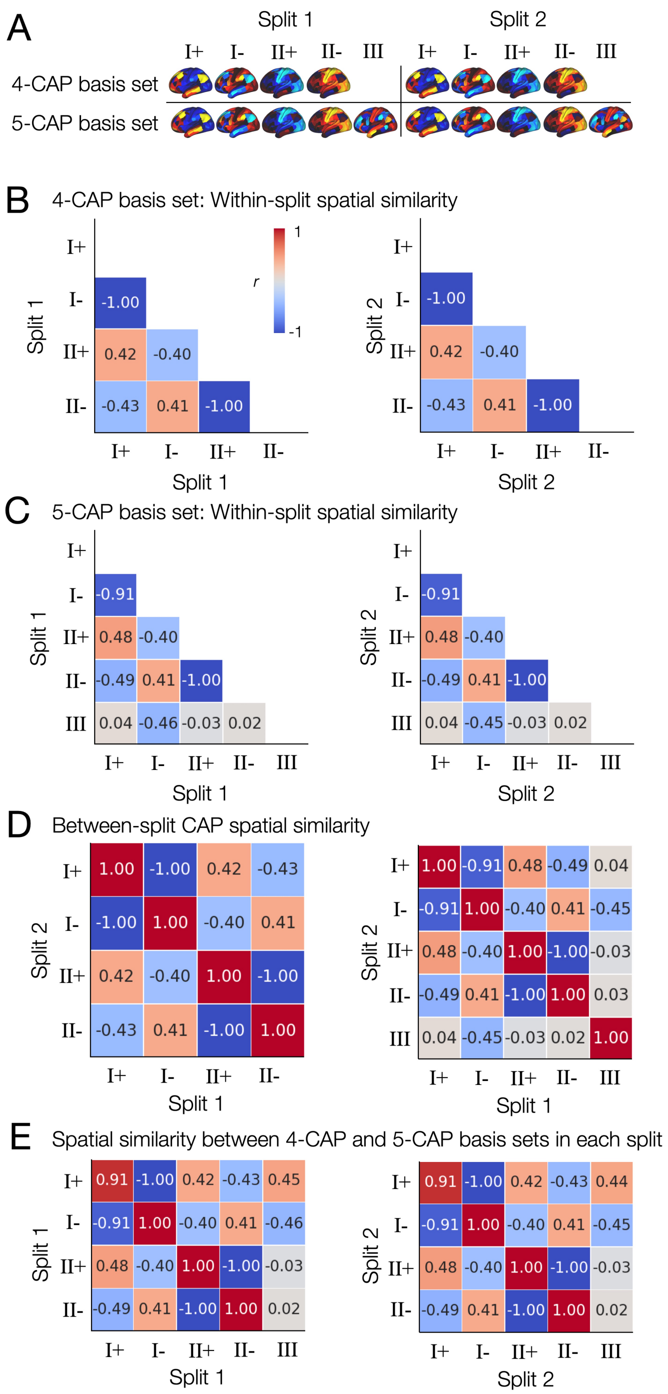


**Fig. S5. Spatial patterns of the basis CAPs are distinct to each other and reproducible using the proposed shuffled split-half analysis. (A)** Spatial patterns of the basis CAPs in each split-half data. The 4-CAP basis set and the 5-CAP basis set were generated independently from the same split-half data, using the hierarchical clustering across 1,000 shuffled split-half resampling, as described in **Fig. S2**. **(B)** Spatial similarity ($r$, correlation coefficient) of the 4-CAP basis set within the split 1 data (left) and within the split 2 data (right). $r$ values were rounded to the nearest 2 decimal digits for visualization. **(C)** Spatial similarity of the 5-CAP basis set within the split 1 data (left) and within the split 2 data (right). **(D)** Spatial similarity of the 4-CAP basis set between the split 1 and 2 data (left) and of the 5-CAP basis set between the split 1 and 2 data (right). **(E)** Spatial similarity between the 4-CAP basis set and the 5-CAP basis set within the split 1 data (left) and within the split 2 data (right).

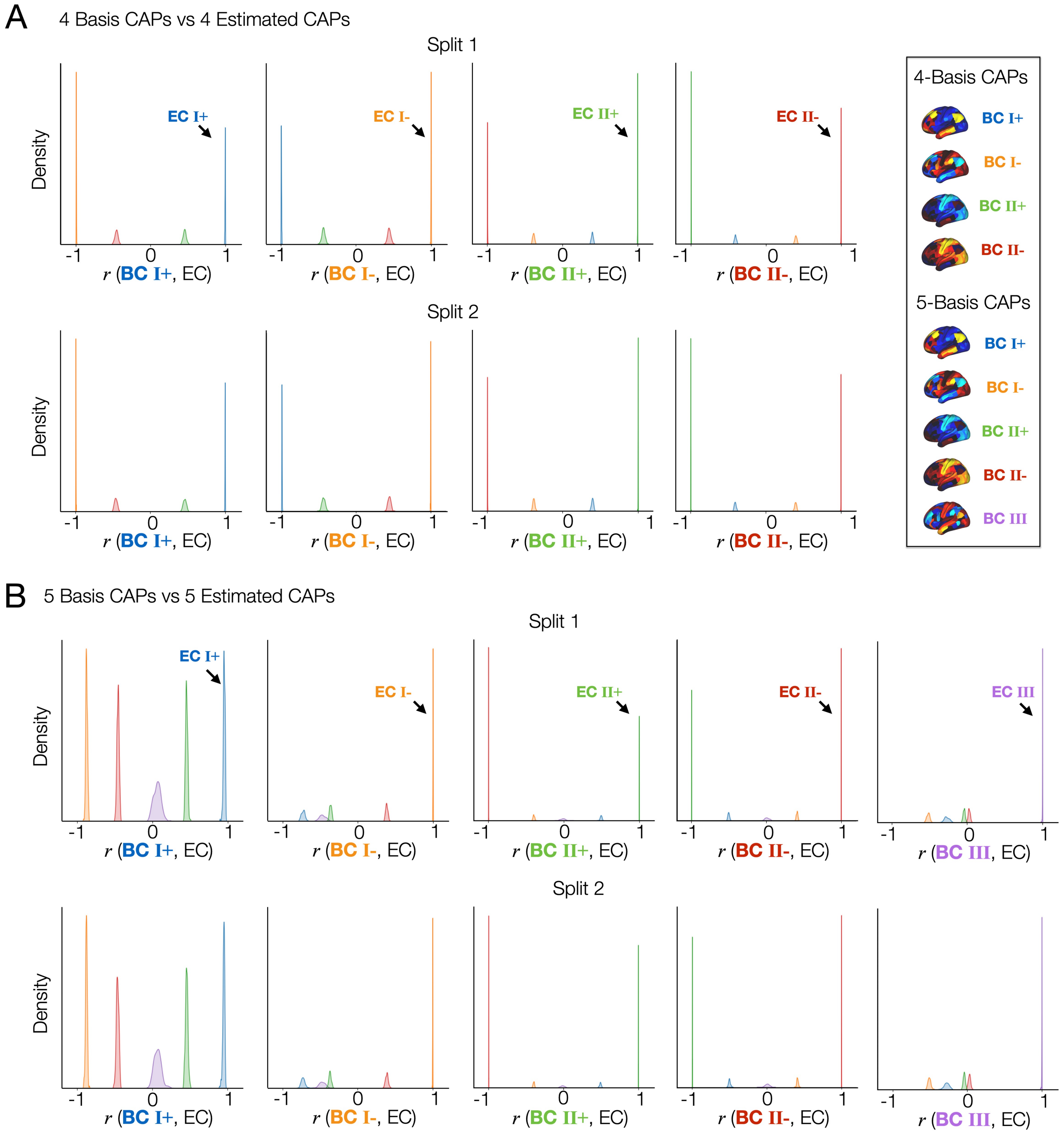


**Fig. S6. Spatial patterns of the CAPs estimated from both splits are reproducible and strongly correlated with at least one of the basis CAPs. (A)** From left to right, the marginal distributions of $r$ between all estimated CAPs (ECs) and each basis CAP (BC) from the 4-CAP basis set are illustrated using kernel density estimation. Results were obtained from the split 1 data (top) and the split 2 data (bottom). Each $r$ value is color-coded using a sorting algorithm to label the corresponding EC using the maximum spatial correlation with BCs. **(B)** From left to right, the marginal distributions of $r$ between all estimated CAPs and each BC from the 5-CAP basis set are illustrated using kernel density estimation.

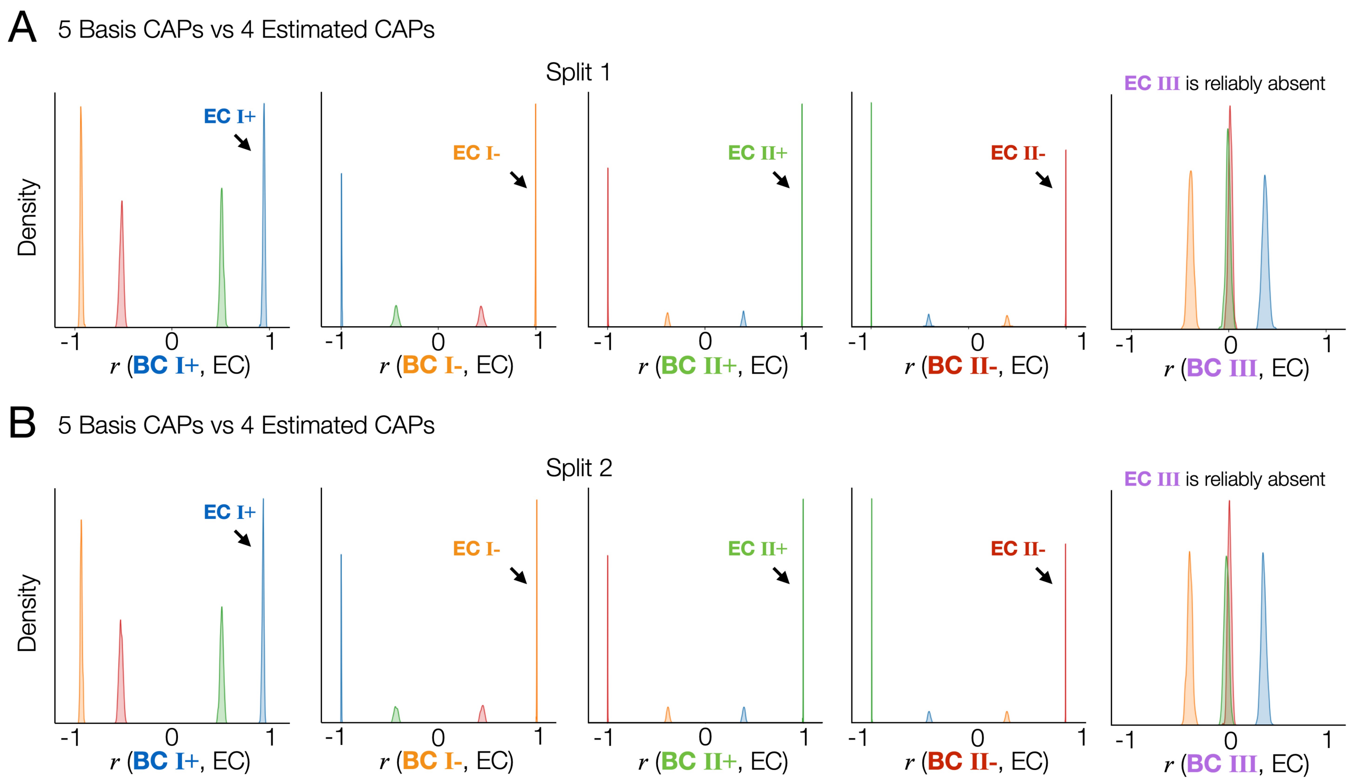


**Fig. S7. The spatial topography of CAP state III is reproducible when it is found in one split and not in another across permutations.**

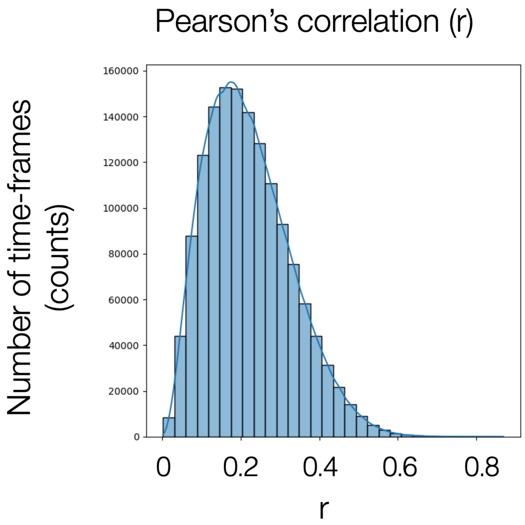


**Fig. S8. The distribution of correlations between individual fMRI time-frames and the estimated basis CAPs (cluster centroid), to which individual time-frames were assigned by K-means clustering.**

DRAFT

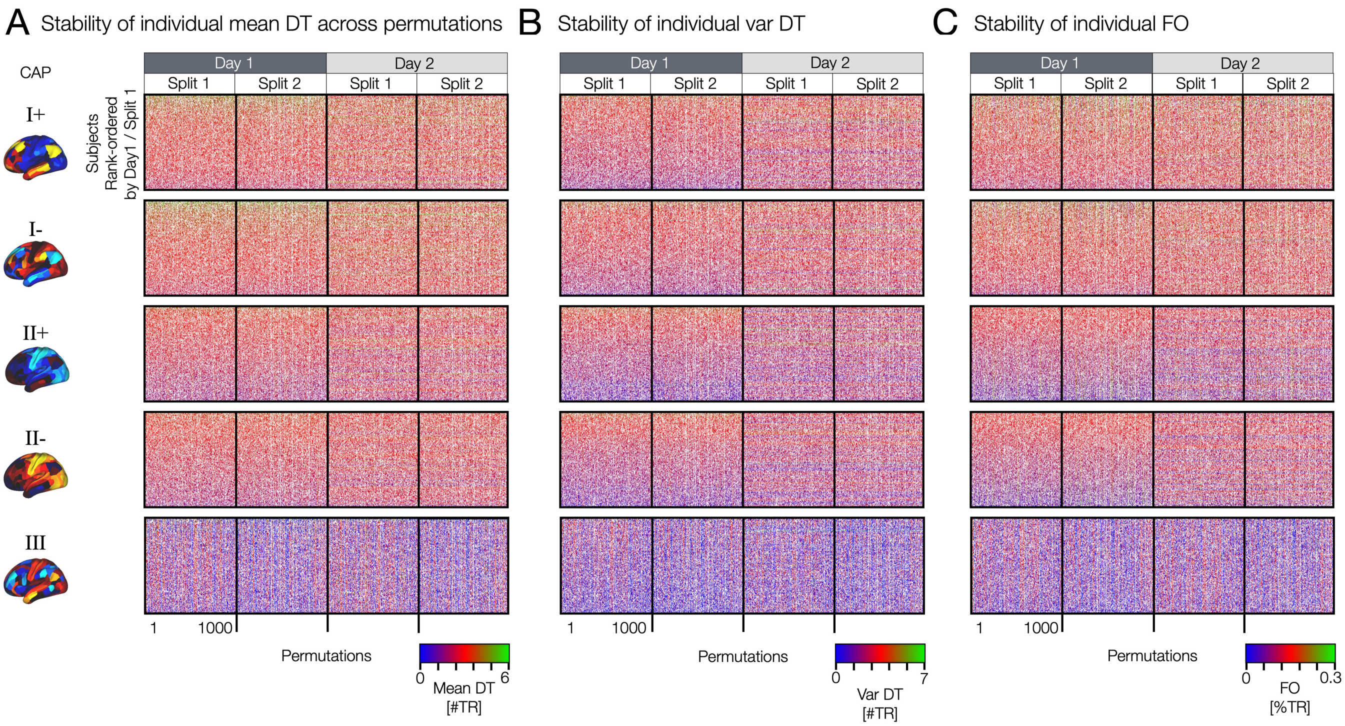


**Fig. S9. Stability of individual mean DT, var DT and FO across permutations**

DRAFT

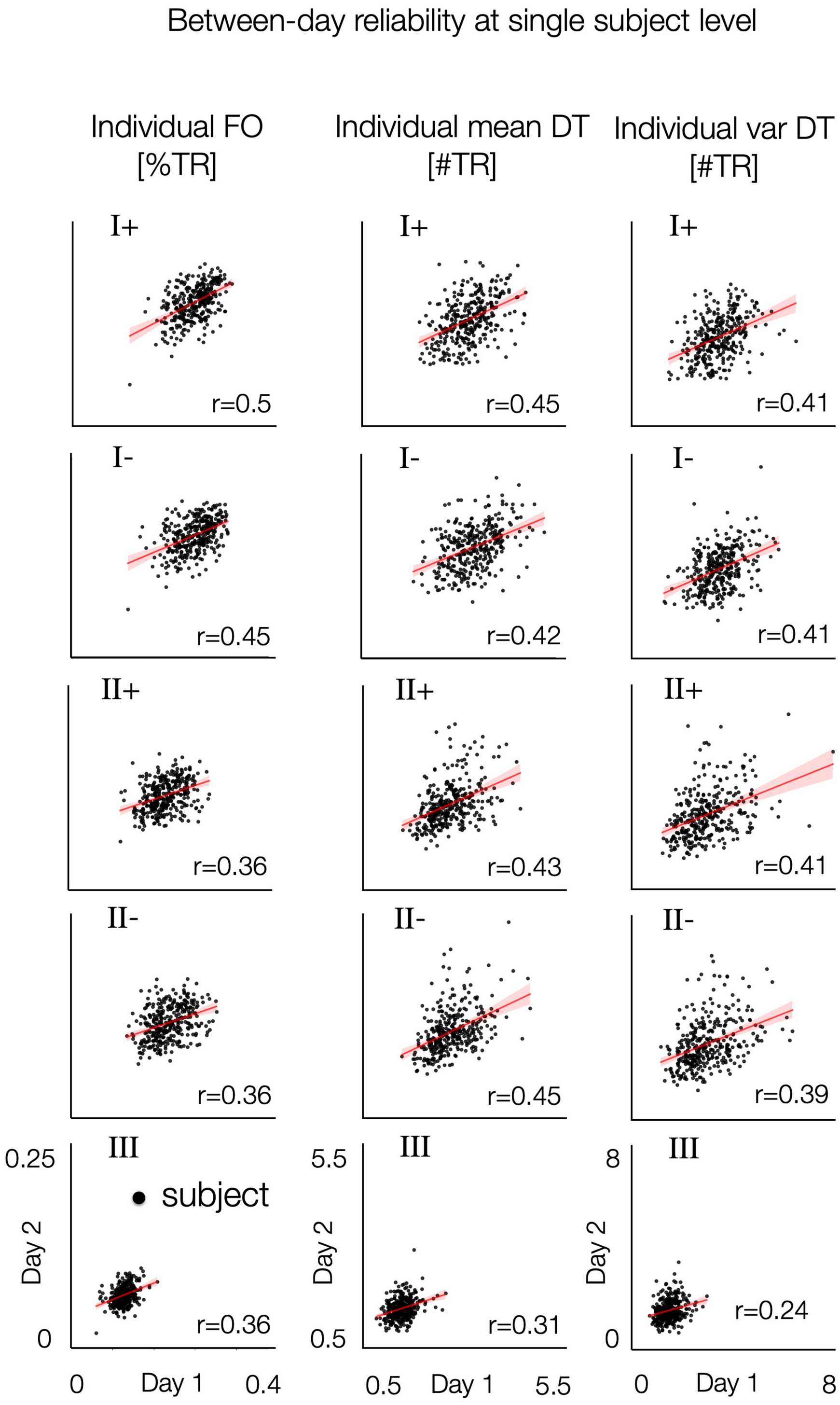


**Fig. S10. Between-day reliability of neural measures at single subject level. Each datapoint in the scatter plot is a subject. For each subject, neural measures were averaged across permutations.**

Similar state-trait features between positive and negative co-activations

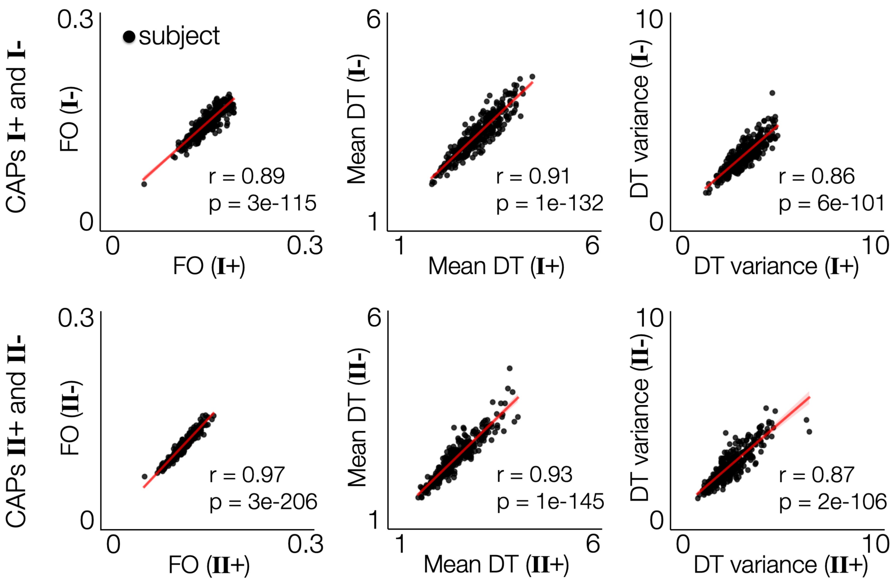


**Fig. S11. Similarity of temporal organizations between positive and negative co-activation patterns.** CAP states I and II have similar FO, mean DT and DT variance across the positive and negative co-activation states (I+ vs I- and II+ vs II-). Each data-point indicate a subject. The temporal metric values across all permutations and two days were averaged within each subject.

DRAFT

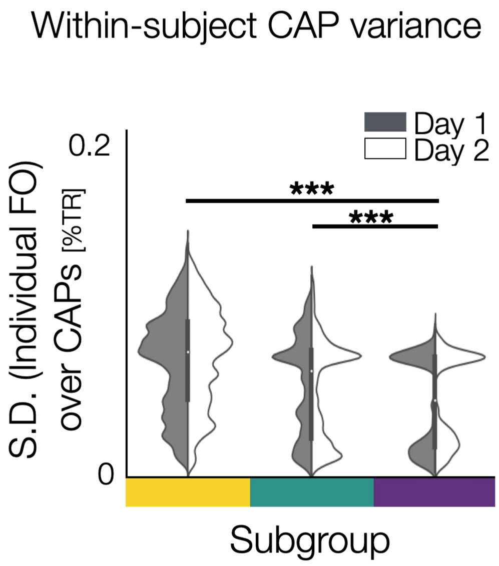


**Fig. S12. Within-subject variance of FO across 5 CAPs across permutations.**

DRAFT

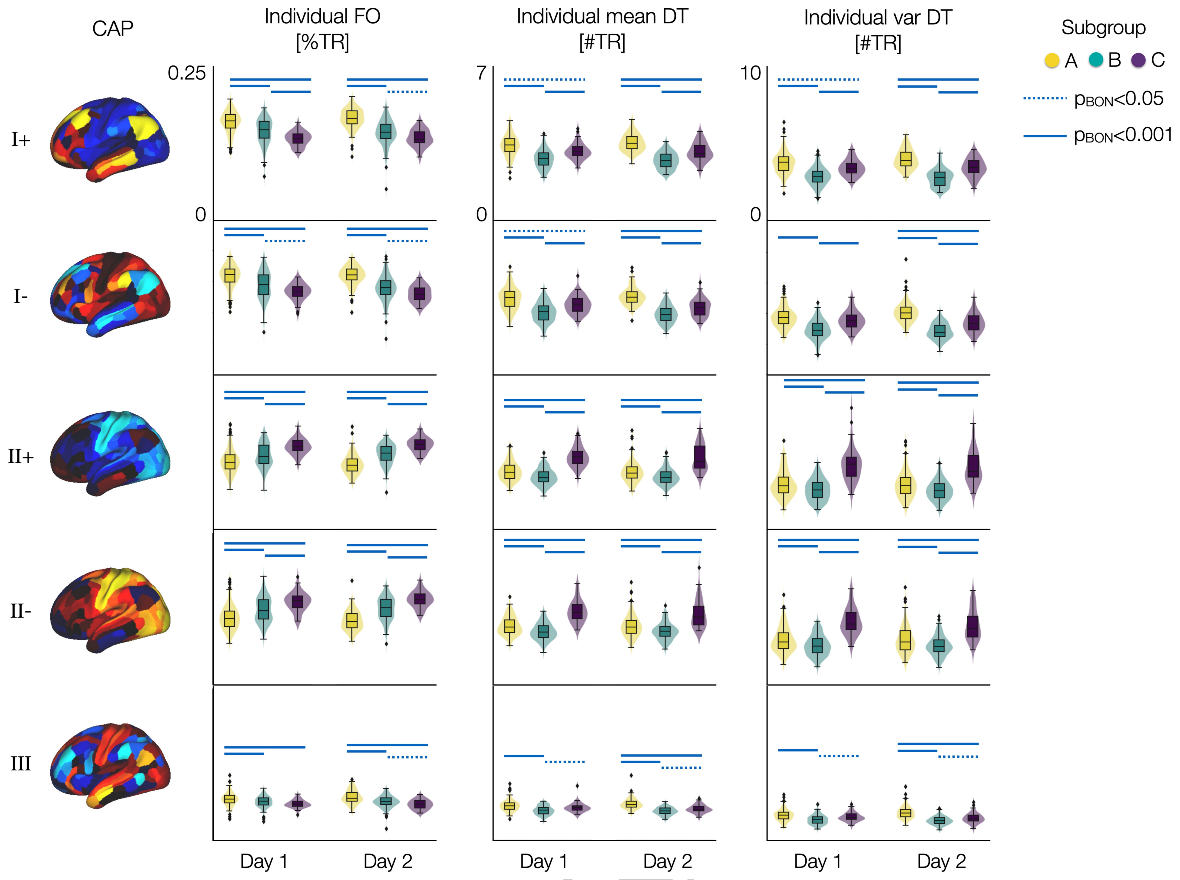


**Fig. S13. Distribution of individual neural measures of spatio-temporal CAP dynamics differ between subgroups.** The distributions of individual FO, mean DT, and var DT of each CAP state are color-coded by the three subgroups. Results from days 1 and 2 data are shown separately and compared between groups. Each data-point indicates a subject. Blue lines: $p$-values with Bonferroni-correction across five CAPs are estimated using two-sided two-sample $t$-tests between groups, $p_{BON} < .001$ (bold) and $p_{BON} < .05$ (dotted).

DRAFT

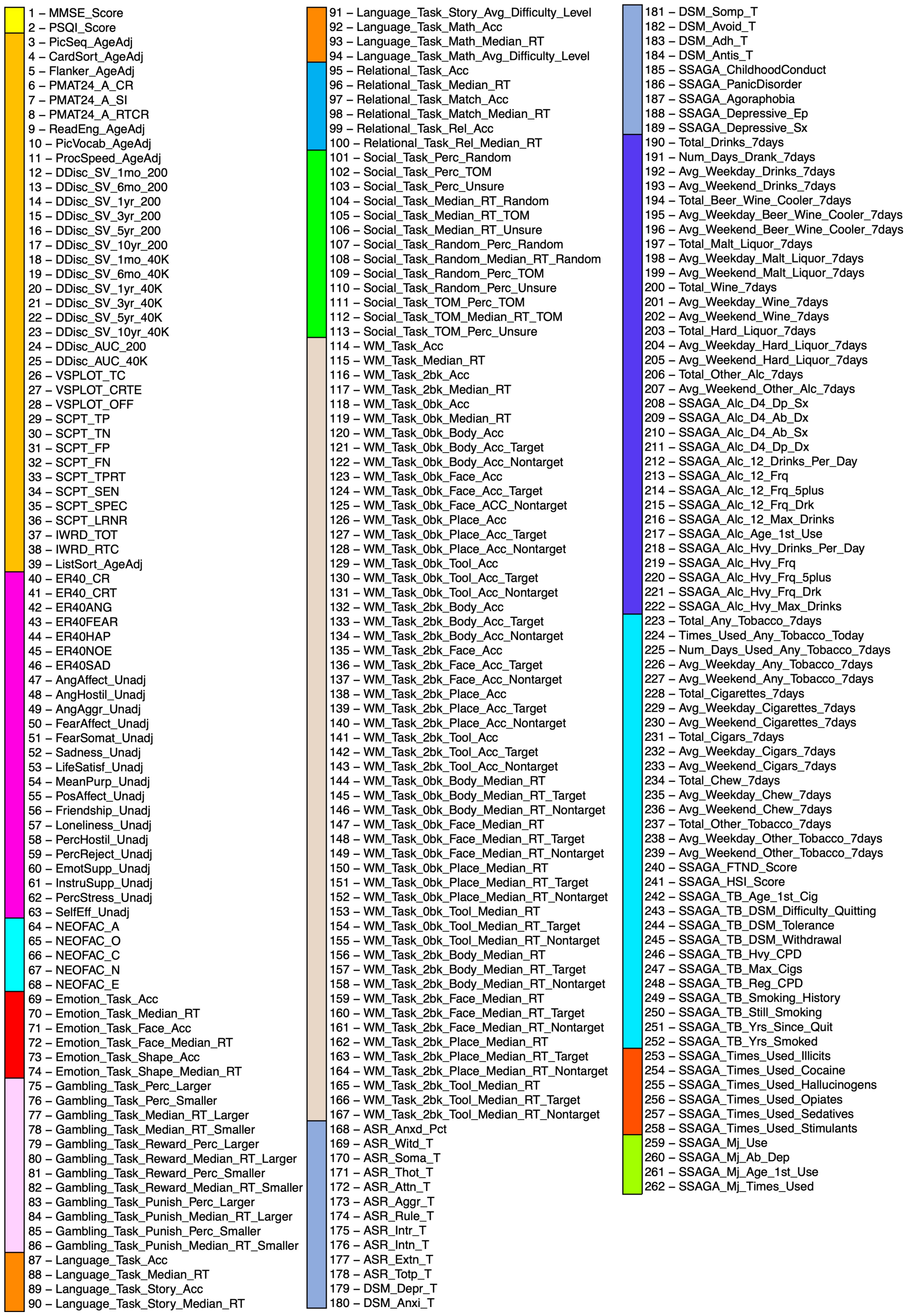


**Fig. S14. List of behavioral variables.** Behavioral variable names are identical to the variable names provided by the HCP data dictionary for the S1200 data release: `HCP_S1200_DataDictionary_April_20_2018.csv`. Check `https://wiki.humanconnectome.org/display/PublicData/HCP-YA+Data+Dictionary-+Updated+for+the+1200+Subject+Release` for details.

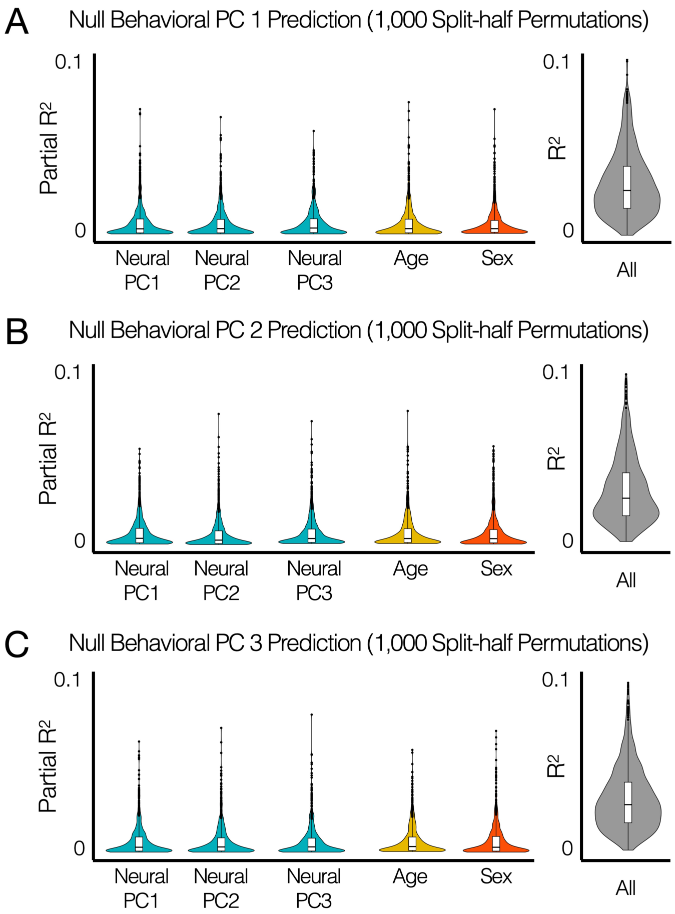


**Fig. S15.** Null data were generated by shuffling individual subjects in behavioral data. Null distributions of partial $R^2$ were estimated for each predictor in the neuro-behavioral association model trained from a split data across 1,000 split-half permutations.

DRAFT

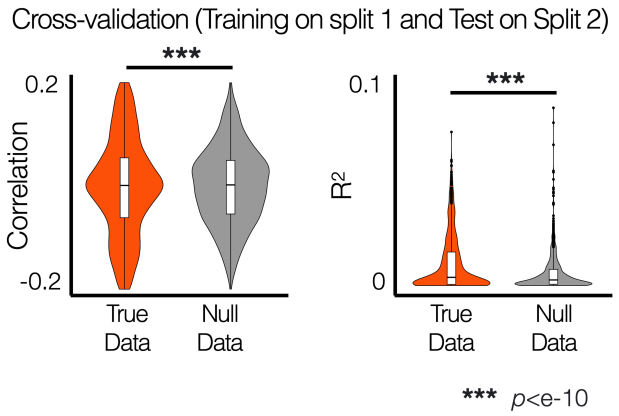


**Fig. S16. Split-half permutation based cross-validation of the prediction model of predicting behavioral PC 1 from neural PCs. The multiple linear regression models were trained using split 1 data and tested on split 2 data in each permutation. Null data were generated by shuffling individual subjects in behavioral data.**

DRAFT

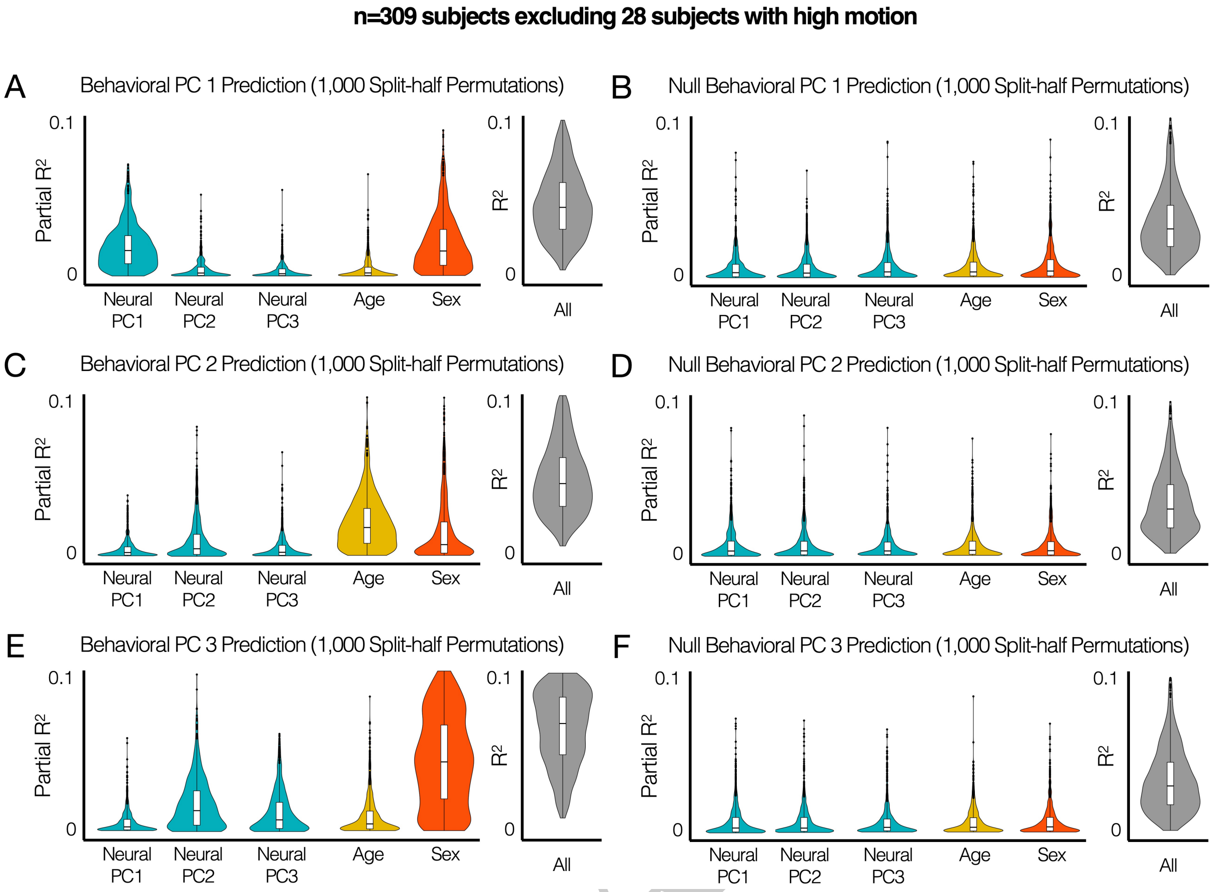


**Fig. S17. Repeating the analysis of neuro-behavioral association excluding subjects with high motion did not change the results.** Among 337 subjects, 28 subjects with excessive motion ($FD > 0.5mm$) were excluded. Across 1,000 permutations, a split of subjects ($n = 154$) was randomly selected, and PCA was performed on neural measures from these subjects. Multiple linear regression models for predicting behavioral PCs from these subjects were estimated. Null data were generated by shuffling individual subjects in behavioral data.
